# The Mechanosensitivity of the Regenerative Response in Bone is Retained with Aging in a Mouse Model of Premature Aging

**DOI:** 10.64898/2026.01.29.702538

**Authors:** Neashan Mathavan, Francisco Correia Marques, Graeme Paul, Sara Lindenmann, Stefanie Wissmann, Dilara Yilmaz, Gisela A Kuhn, Esther Wehrle, Ralph Müller

## Abstract

Aging impairs the regenerative capacity of bone and is associated with poor healing outcomes. The mechanical environment is of fundamental importance to the regenerative response in bone – yet the effect of aging on the mechano-responsive capacity of bone regeneration remains largely unresolved. To investigate age-dependent mechanobiological responses in bone regeneration, we established an experimental framework consisting of: (i) an established femur defect mouse model, (ii) the use of PolgA^D257A/D257A^ (PolgA) mice - a mouse model of premature aging, and (iii) our recently established spatial transcriptomics–based “mechanomics” platform which permits gene expression to be analyzed as a function of the local *in vivo* mechanical environment. Aging impaired the regenerative response in PolgA mice, resulting in an increased occurrence of delayed and non-unions, delayed bone formation / resorption responses, impaired osteogenesis and delayed mineralization of new bone. Cyclic mechanical loading significantly enhanced the regenerative response in young PolgA mice inducing sustained bone formation, suppressing bone resorption, and enhancing mineralization, with the strongest effects observed in peripheral regions of the fracture site. In aged PolgA mice, the mechanosensitivity of the regenerative response was retained with an anabolic response localized to the defect center. Cyclic mechanical loading applied during the reparative and remodelling phases of fracture healing thus represents a potential translational strategy to harness the mechanosensitivity of aged bone.

## Introduction

Aging is associated with the dysregulation of mechanobiological mechanisms that impairs the ability of cells to sense and adapt to extracellular mechanical stimuli^1^. The mechanical environment is of critical importance to the regenerative response in bone. However, no consensus exists on the effect of aging on the mechanosensitivity of bone regeneration. Certain characteristics of bone aging have been identified with the potential to influence bone mechanosensitivity and its subsequent regenerative capacity. These include: decreased osteocyte density^2,3^, reducing the primary mechanosensing cell population; deterioration of the lacuno-canalicular network (LCN)^4– 6^, disrupting mechanotransduction pathways; and declining osteogenic cell populations^7^ coupled with compromised angiogenesis^8^, limiting the tissue’s ability to respond appropriately to mechanical cues^9^. Given the higher incidence of recalcitrant fractures in elderly populations, insights into the mechanisms by which aging alters bone mechanosensitivity may provide much-needed novel therapeutic strategies to enhance bone regeneration in these patients.

Investigating the influence of aging on the mechanosensitivity of bone healing presents multifaceted challenges. Although rodent models are well-established within the field, the use of naturally aged animals requires considerably longer investigation timelines^10^, and maintaining such colonies is resource intensive. Furthermore, *in vivo* mechanical loading applied at the organ scale is heterogeneously distributed throughout fracture sites, resulting in complex mechanical environments with distinct regions of high and low strain^11^. Consequently, each cell within a fracture site resides within different local mechanical microenvironments. This heterogeneity in local mechanical environments renders measuring cellular responses to mechanical stimuli within bone tissue technically challenging.

To address these challenges, we have developed an integrative experimental framework to investigate age-dependent mechanobiological responses in fracture healing. At the core of our framework is an established femur defect mouse model consisting of mid-diaphyseal femoral defects stabilized with an external fixator^12^. Our model permits longitudinal *in vivo* micro-computed tomography (micro-CT) imaging and cyclic mechanical loading of the fracture site via the external fixator. We recently demonstrated a cyclic mechanical loading protocol using our femur defect mouse model that significantly enhanced osteogenic responses during fracture healing in young adult mice^11–13^. To investigate the effects of aging on the regenerative capacity and mechanosensitivity of bone, we have incorporated PolgA^D257A/D257A^ (PolgA) mice – a mouse model of premature aging^14^ within our framework. PolgA mice present a premature aging phenotype due to the accumulation of mitochondrial DNA (mtDNA) point mutations at rates 3 – 5 fold higher compared to wild type mice^15^. In our characterizations of their musculoskeletal phenotype, PolgA mice were found to exhibit the premature onset of clinically-relevant musculoskeletal aging characteristics including frailty, osteosarcopenia, and degeneration of the LCN^5,16–18^. To link mechanical loading applied at the organ-scale to molecular responses, we incorporated our recently established spatial transcriptomics–based “mechanomics” platform within our experimental framework, permitting gene expression to be analyzed as a function of the mechanical environment^5^. Combined, this experimental framework provides a unique opportunity to dissect how aging affects the mechanosensitivty of bone regeneration across multiple length scales: from tissue architecture down to cellular and molecular mechanisms.

Emerging pre-clinical research in the treatment of postmenopausal osteoporosis suggests that combining load-bearing physical activity with drug therapies yields superior outcomes in bone strength compared to either approach alone^19,20^. With these findings as a basis, we hypothesized that cyclic mechanical loading would enhance the regenerative response at fracture sites in aged bone by augmenting the intrinsic anabolic response. Accordingly, our aim was to determine whether cyclic mechanical loading applied during the reparative and remodelling phases of fracture healing represents a potential translational strategy to harness the mechanosensitivity of aged bone.

## Results

### Aging impairs the regenerative response in PolgA mice

Fracture sites in 12-week-old “Young” and 35-week-old “Aged” PolgA mice were classified as unions (if cortical bridging was present by week 3 post-surgery), delayed unions (if cortical bridging was observed between weeks 4 – 7 post-surgery), or non-unions (if no bridging was present up to 7 weeks-post surgery). Classifications were based on previously reported healing outcomes in WT mice using the same model^12^. Consistent with age-related impaired regeneration, an increased incidence of delayed and non-unions was observed in Aged mice. In comparisons between fracture sites which were classified as unions, the regenerative response varied both spatially and temporally between Young and Aged PolgA mice. Sites of bone formation, quiescence and resorption at the fracture site using time-lapsed *in vivo* micro-CT imaging are presented at weekly intervals in Figure 2.

**Fig. 1.**
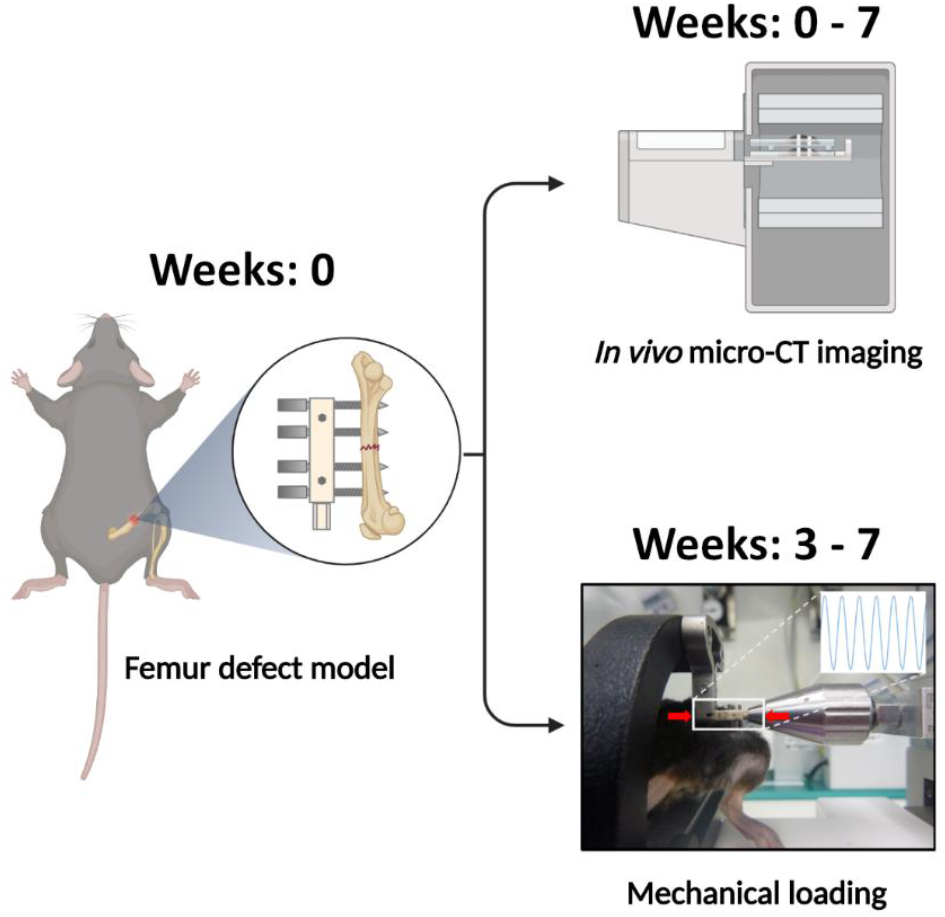
Overview of experiment design. At Week 0, mid-diaphyseal femoral osteotomies are introduced in the right femur of mice and stabilized with an external fixator. Time-lapsed *in vivo* micro-CT imaging is performed weekly at the fracture site (weeks 0-7; 10.5 μm nominal resolution). Mice which exhibit bridging at 3 weeks post-surgery are subdivided into Loaded and Control groups. At weeks 3 – 7, mice received individualized cyclic loading (up to 16 N) or 0 N sham-loading three times per week. All mice are euthanized at 7 weeks post-surgery. Illustration adapted from Mathavan et al.^11^.

**Fig. 2.**
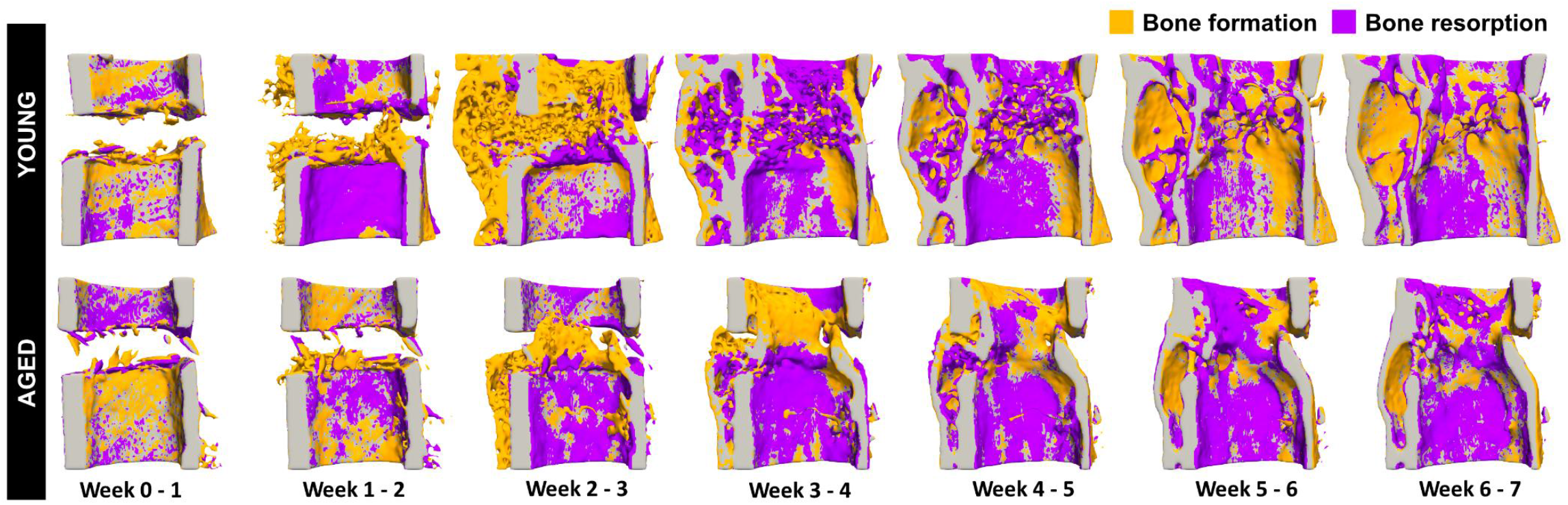
Visualization of sites of bone formation, quiescence, and resorption in Young and Aged PolgA mice. Sites of bone formation (orange) and bone resorption (purple) are identified via registration of time-lapsed *in vivo* micro-CT images (threshold: 395 mg hA/cm3, voxel size = 10.5 μm). Representative samples per group were identified based on BV/TV at week 7 in fractures classified as unions. Visualization performed using Paraview (version 5.7.0).

In our quantitative morphometric analyses of time-lapsed *in vivo* micro-CT imaging data (presented as mean ± SD), aging was associated with a delayed and diminished regenerative response. In assessments of fracture healing progression in young vs. aged mice, modified Radiographic Union Scores (mRUST) at three weeks post-osteotomy were significantly lower for aged mice compared to young mice (7.8 ±2.0% vs. 10.1 ±1.3%; p ≤ 0.01). Bone volume fraction (BV/TV) in the defect center (DC) of Young mice was significantly higher compared to Aged mice at week 2 (27.3 ±7.3% vs. 11.4 ±1.8%; p ≤ 0.01) and week 3 (46.9 ±8.7% vs. 25.8 ±8.4%; p ≤ 0.01) (Fig 3. [A]). Similarly in peripheral regions, BV/TV was significantly higher at week 3 in the defect periphery (DP) (27.4 ±9.6% vs. 11.0 ±8.4%; p < 0.05) (Fig 3. [B]) and week 2 in the cortex periphery (FP) (26.8 ±8.6% vs. 13.9 ±4.6%; p < 0.05) (Fig 3. [D]). However, BV/TV values comparable to Young mice were achieved in both DC and DP regions of Aged mice from week 4 onwards suggestive of a delayed response with aging (Fig 3. [A][B]). In contrast, BV/TV values remained significantly lower in the FP region of Aged mice from week 4 onwards reflecting a diminished osteogenic response in peripheral regions (Fig 3. [D]; p < 0.05).

**Fig. 3.**
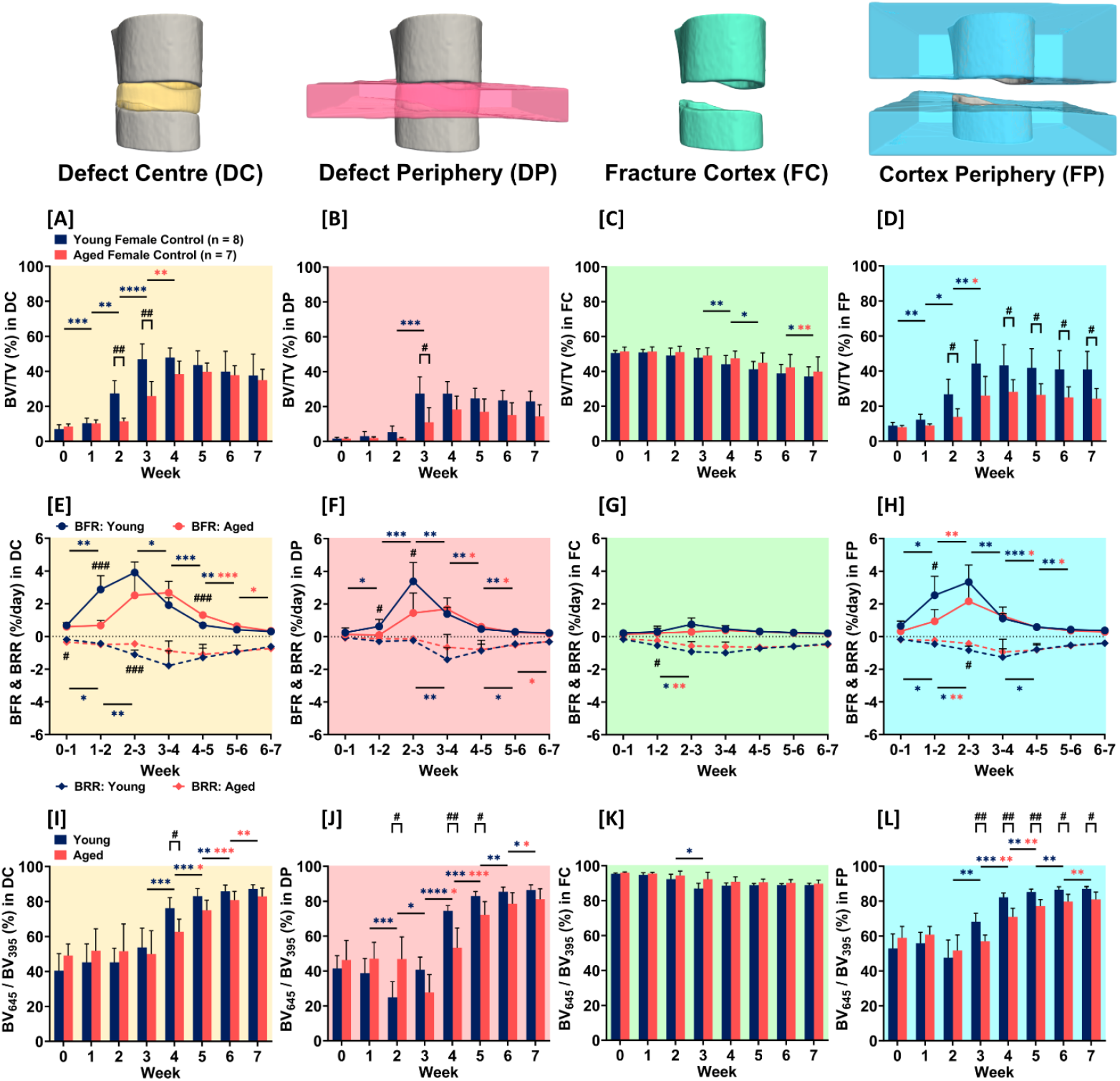
Regenerative response in Young vs. Aged Female mice. Quantitative morphometric analyses were performed in four volumes of interest: the defect center (DC), the defect periphery (DP), the existing fracture cortices together with the medullary cavity (FC), and the cortex periphery (FP). Three parameters are presented: **[A]-[D]** BV/TV where BV is the bone volume in each VOI normalized to TV (DC for DC and DP, FC for FC and FP), **[E]-[H]** bone remodelling rates: bone formation rate (BFR – solid line) and bone resorption rate (BRR – dashed line), and **[I]-[L]** BV_645_/BV_395_ representing the degree of bone mineralization given as the ratio of bone volume with a density ≥ 645 mg HA/cm^3^to the total osseous volume (threshold ≥ 395 mg HA/cm^3^). Young mice: n = 8, Aged mice: n = 7. Significance criteria: **#** represents comparisons between Young and Aged mice at each timepoint (# p < 0.05, ## p < 0.01, ### p < 0.001, #### p < 0.0001), **\*** represents comparisons within groups between timepoints (* p < 0.05, ** p < 0.01, *** p < 0.001, **** p < 0.0001).

Analysis of bone formation and bone resorption rates (BFR and BRR) at the fracture sites of Young and Aged mice further corroborated these findings. In DC regions, BFR in both groups were characterized by a rapid rise that plateaued and subsequently declined; however, this sharp increase in BFR occurred between weeks 1 – 2 in Young mice, whereas the corresponding increase in Aged mice occurred between weeks 2 – 3 (Fig 3. [E]). In DP regions, the formation response in Aged mice was diminished compared to Young mice between weeks 1 – 2 (0.1 ±0.04 % per day vs. 0.6 ±0.4 % per day; p ≤ 0.05) and between weeks 2 – 3 (1.5 ±1.2 % per day vs. 3.4 ±1.2 % per day; p ≤ 0.05) (Fig 3. [F]). Similarly, the formation response in FP regions was significantly lower in Aged mice between weeks 1 – 2 (0.9 ±0.7 % per day vs. 2.5 ±1.2 % per day; p ≤ 0.05) (Fig 3. [H]). Consistent with the coupling of bone formation with bone resorption, BRRs tended to be lower in Aged mice compared to Young mice across all regions (Fig 3. [E][F][G][H]) – with statistically significant differences in BRR observed at early time points (Fig 3. [E][G][H]).

Site specific differences were observed in comparisons of the mineralization of the fracture sites of Young and Aged mice, with Aged mice exhibiting a consistent one-week delay in the onset of mineralization (Fig 3. [I][J][L]). In DC regions, both Young and Aged mice exhibited comparable degrees of mineralization between weeks 0 – 3 (Fig 3. [I]). Progressive increases in mineralization were subsequently observed in Young mice between weeks 3 – 6 (53.7 ±11.1% at week 3 vs. 85.8 ±3.6% at week 06) (Fig 3. [I]) and in Aged mice between weeks 4 – 7 (62.6 ±7.3% at week 4 vs. 82.9 ±4.8% at week 07) (Fig 3. [I]). Similarly in FP regions, increases in mineralization were observed in Young mice from week 2 onwards, whereas corresponding increases in Aged mice were present from week 3 onwards (Fig 3. [L]). In DP regions, the fraction of highly mineralized tissue declined sharply at week 2 in Young mice and week 3 in Aged mice, reflecting rapid deposition of new, lowly mineralized bone (Fig 3. [J]). Following this decline, progressive increases across all subsequent timepoints were observed, indicating maturation and mineralization of the newly formed bone.

Collectively, quantitative micro-CT morphometric analyses demonstrate that aging impairs the regenerative response in PolgA mice, characterized by an increased incidence of delayed and non-unions, delayed bone formation / resorption responses in central regions (DC), diminished osteogenic responses in peripheral sites (DP, FP), and delayed maturation and mineralization of newly formed bone.

### Cyclic mechanical loading significantly enhances the regenerative response in Young PolgA mice

No significant differences in bone morphometric parameters were observed between Control and Loaded groups in the pre-loading healing period (week 0 to week 3) across all defect VOIs (Fig 4. [A]; Fig 5.). Cyclic mechanical loading of the fracture sites of young mice (between week 3 to week 7), significantly enhanced the regenerative response (Fig 4. [A]) – as demonstrated in the contrast in ΔBV/TV between Loaded mice (DC: +35.6%, DP: +104.2%, FC: +11.6%, FP: +100.2%) vs. Control mice (DC: -19.9%, DP: -16.4%, FC: -22.5%, FP: -7.8%) (Fig 5.).

**Fig. 4.**
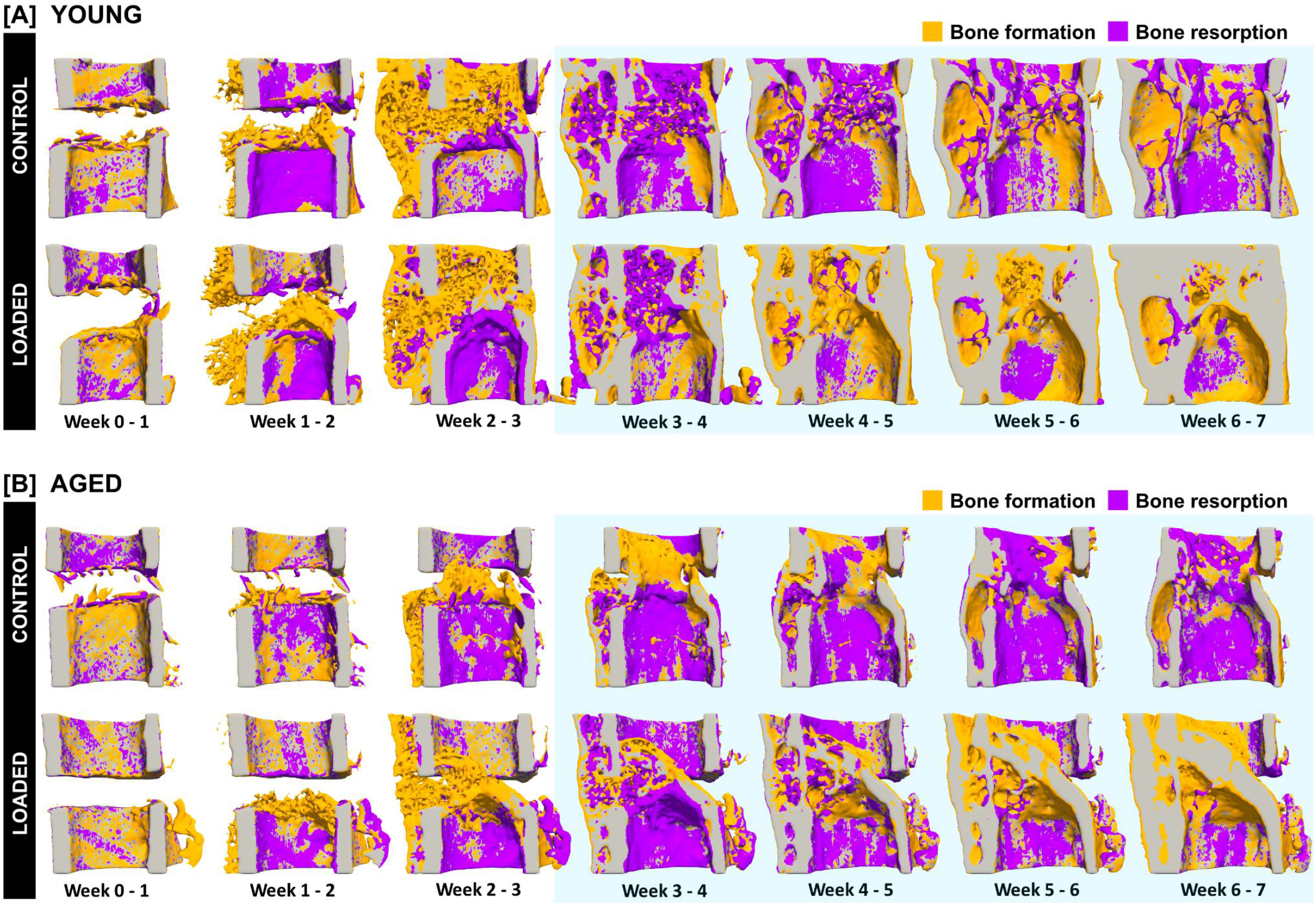
Visualization of sites of bone formation, quiescence, and resorption in Control and Loaded mice in [A] Young and [B] Aged PolgA mice. Sites of bone formation (orange) and bone resorption (purple) are identified via registration of time-lapsed *in vivo* micro-CT images (threshold: 395 mg hA/cm3, voxel size = 10.5 μm). Representative samples per group were identified based on BV/TV at week 7 in fractures classified as unions. Shaded regions (blue) correspond to timepoints at which mechanical loading was applied. Visualization performed using Paraview (version 5.7.0).

**Fig. 5.**
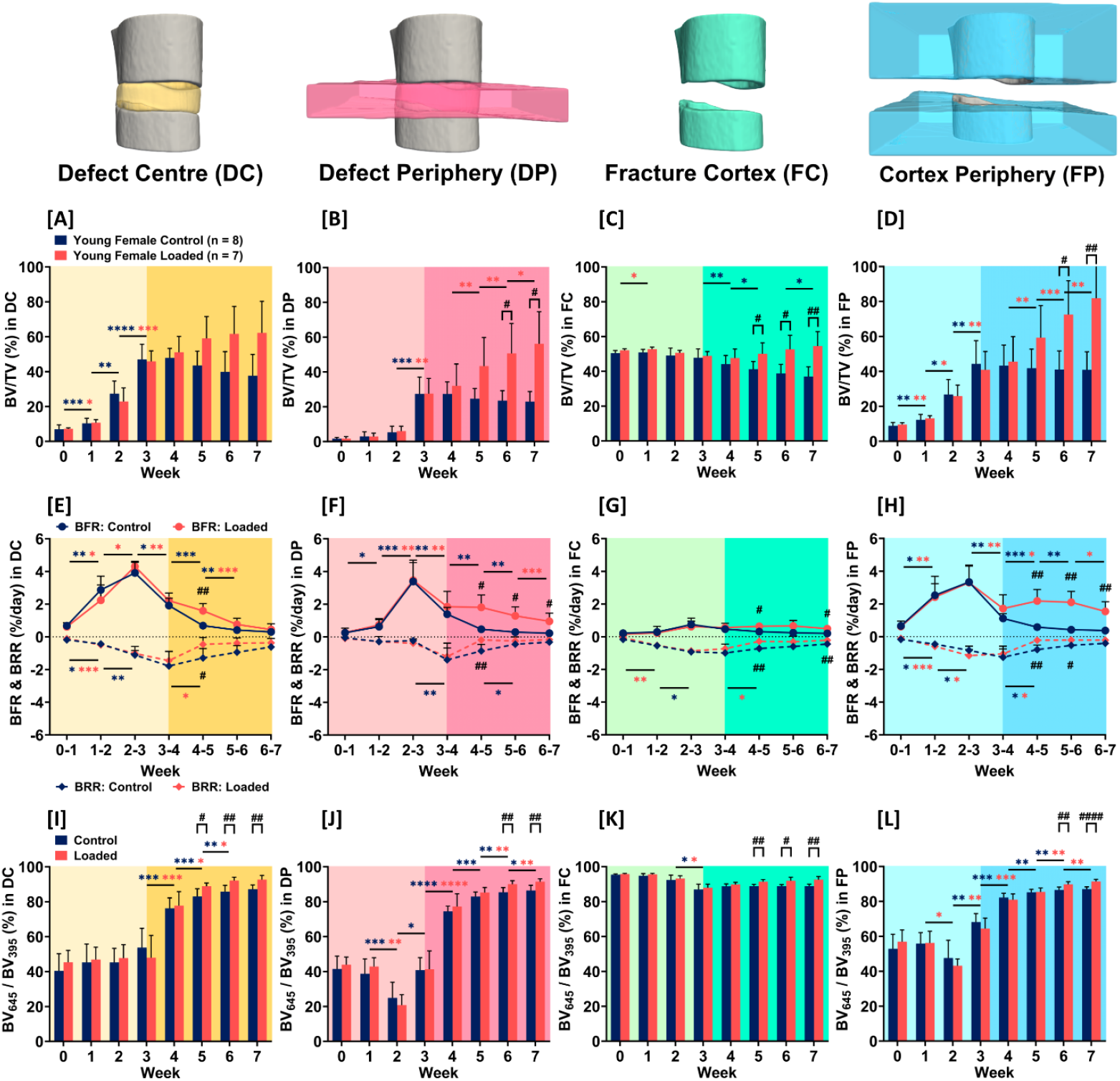
Regenerative response in Young Control vs. Loaded Female mice. Quantitative morphometric analyses were performed in four volumes of interest: the defect center (DC), the defect periphery (DP), the existing fracture cortices together with the medullary cavity (FC), and the cortex periphery (FP). Shaded regions in each plot correspond to time points at which loading was applied 3x per week. Three parameters are presented: **[A]-[D]** BV/TV, **[E]-[H]** bone remodelling rates: bone formation rate (BFR – solid line) and bone resorption rate (BRR – dashed line), and **[I]-[L]** BV_645_/BV_395_ representing the degree of bone mineralization. Control mice: n = 8, Loaded mice: n = 7. Significance criteria: # represents comparisons between Young and Aged mice at each timepoint (# p < 0.05, ## p < 0.01, ### p < 0.001, #### p < 0.0001), * represents comparisons within groups between timepoints (* p < 0.05, ** p < 0.01, *** p < 0.001, **** p < 0.0001).

The response to cyclic mechanical loading was site-specific and predominantly localized to peripheral regions. BV/TV in DC regions diverged between Control and Loaded groups from week 3 onwards (Fig 5. [A]). In Control mice, a trend of progressive decreases in the parameter was present. Conversely, in Loaded mice, a trend of progressive increases was observed (Fig 5. [A]). At week 7, BV/TV in the DC region was 37.6 ± 12.2% in Control mice versus 62.2 ± 18.1% in Loaded mice (ns).

In peripheral regions (DP, FP), mechanical loading resulted in significant increases in BV/TV at week 6 (DP, FP: p ≤ 0.05) and week 7 (DP: p ≤ 0.05; FP: p ≤ 0.01) (Fig 5. [B][D]). In both peripheral regions, BV/TV significantly increased between weeks 4 – 5 (DP, FP: p ≤ 0.01), weeks 5 – 6 (DP: p ≤ 0.01, FP: p ≤ 0.001) and weeks 6 – 7 (DP: p ≤ 0.05, FP: p ≤ 0.01) in Loaded mice, whereas no significant changes were observed in Control mice (Fig 5. [B][D]). Similarly, in the cortical and medullary region (FC), BV/TV was significantly higher in response to mechanical loading at week 5 (p ≤ 0.05), week 6 (p ≤ 0.05) and week 7 (p ≤ 0.01) (Fig 5. [C]). Longitudinally, Control mice exhibited a progressive decline in BV/TV in the FC region, with significant decreases between weeks 3 – 4 (p ≤ 0.01), weeks 4 – 5 (p ≤ 0.05) and weeks 6 – 7 (p ≤ 0.05) (Fig 5. [C]).

Dynamic bone morphometric parameters further corroborated the predominant localization of the response to peripheral regions. In both groups, BFR increased during the early timepoints, peaking between weeks 2 and 3, and then declined over subsequent weeks (Fig 5. [E][F][G][H]). However, the rate of decline differed between Control and Loaded groups. In Loaded mice, this decline was partially attenuated, resulting in a sustained bone formation rate and a diminished bone resorption rate – effects that were most pronounced in peripheral regions (DP, FP) (Fig 5. [F][H]). In DP regions, BFR was significantly higher in Loaded mice compared with Control mice between weeks 4 – 5 (p ≤ 0.05), weeks 5 – 6 (p ≤ 0.05), and weeks 6 – 7 (p ≤ 0.05), whereas BRR was significantly lower in Loaded mice between weeks 4 – 5 (p ≤ 0.01) (Fig 5. [F]). Similarly, in FP regions, BFR was significantly higher in Loaded mice compared with Control mice between weeks 4 – 5 (p ≤ 0.01), weeks 5 – 6 (p ≤ 0.01), and weeks 6 – 7 (p ≤ 0.01), while BRR was significantly reduced in Loaded mice between weeks 4 – 5 (p ≤ 0.01) and weeks 5 – 6 (p ≤ 0.05) (Fig 5. [H]).

Mineralization dynamics across all VOIs were comparable between Control and Loaded groups during the pre-loading healing period (week 0 to week 3) (Fig 5. [I][J][K][L]). Cyclic mechanical loading resulted in a significantly greater fraction of mineralized bone volume across all VOIs in Loaded mice at weeks 6 and 7 compared to Control mice (Fig 5. [I][J][K][L]). These findings are indicative of callus maturation and suggest cyclic mechanical loading promotes rather than delays mineralization.

Site-specific differences in mineralization dynamics were evident in comparisons of Control and Loaded groups. In DC regions, both groups exhibited significant increases in mineralized bone fraction at each successive timepoint between weeks 3 – 6, reflecting continuing callus maturation (Fig 5. [I]). Moreover, loaded mice displayed a significantly greater fraction of mineralized bone compared with Controls at weeks 5 (p ≤ 0.05), 6 (p ≤ 0.01), and 7 (p ≤ 0.01) (Fig 5. [I]). In both peripheral regions (DP, FP), the fraction of highly mineralized bone volume declined between weeks 1 – 2 followed by progressive increases across all subsequent timepoints in both Control and Loaded groups (Fig 5. [J][L]). These findings are indicative of the formation of lowly-mineralized new bone in these regions that subsequently undergoes mineralization. This observation was not present in the DC region, underscoring the site-specific nature of the response to mechanical loading and its predominant localization to peripheral regions in young mice.

Collectively, these findings demonstrate that cyclic mechanical loading significantly enhances the regenerative response in young PolgA mice, inducing sustained bone formation, suppressed bone resorption, and enhanced mineralization, with the strongest effects observed in peripheral regions (DP, FP).

### Mechanosensitivity of the regenerative response is retained in Aged PolgA mice

In Aged mice, the regenerative response to cyclic mechanical loading exhibited distinct spatio-temporal dynamics compared to Young mice. In contrast to Young mice, cyclic mechanical loading induced a robust anabolic response within the DC region (Fig 4. [B]; Fig 6. [A]), with minimal responses in peripheral regions (DP, FP) (Fig 6. [B][D]).

**Fig. 6.**
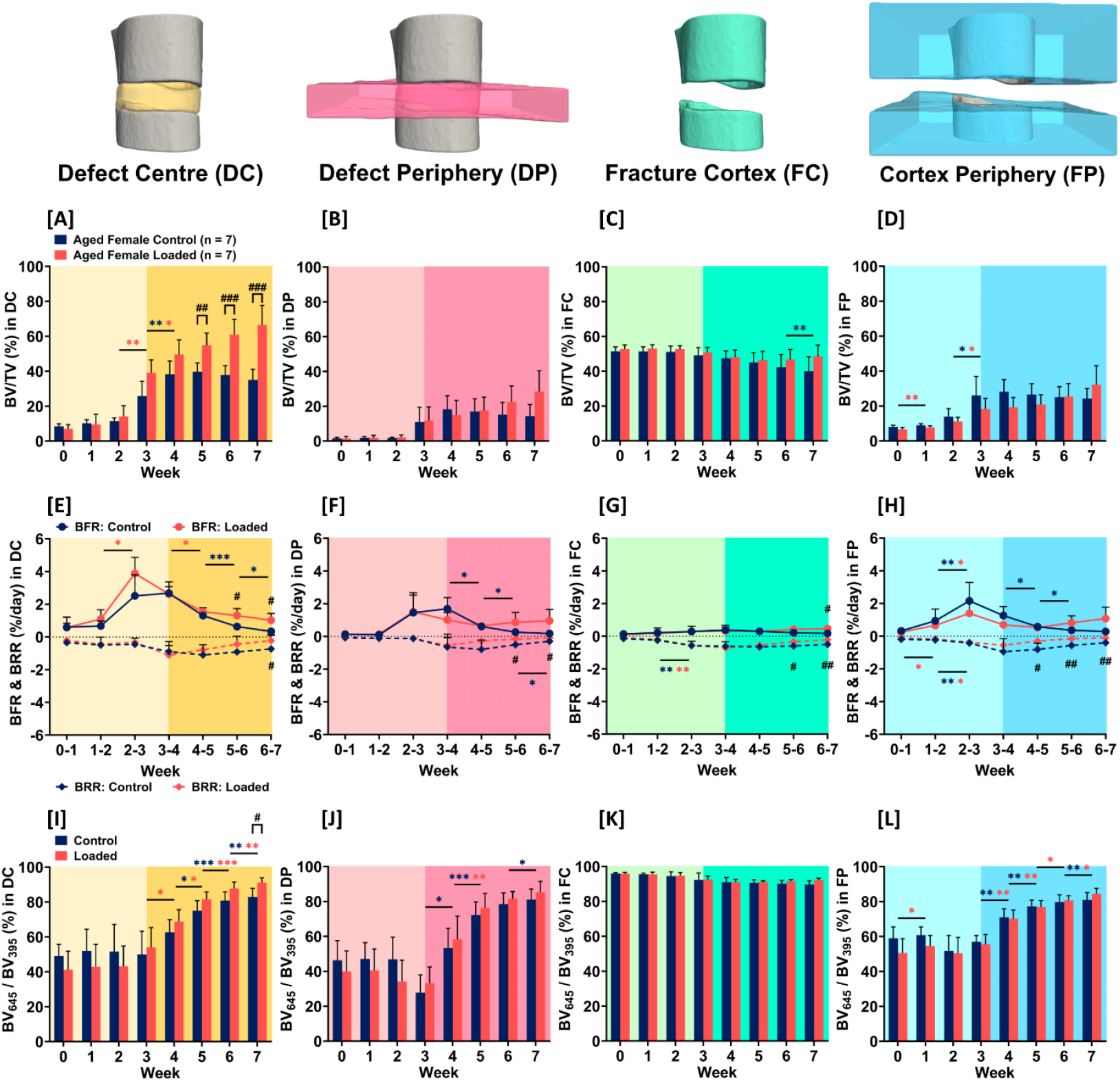
Regenerative response in Aged Control vs. Loaded Female mice. Quantitative morphometric analyses were performed in four volumes of interest: the defect center (DC), the defect periphery (DP), the existing fracture cortices together with the medullary cavity (FC), and the cortex periphery (FP). Shaded regions in each plot correspond to time points at which loading was applied 3x per week. Three parameters are presented: **[A]-[D]** BV/TV, **[E]-[H]** bone remodelling rates: bone formation rate (BFR – solid line) and bone resorption rate (BRR – dashed line), and **[I]-[L]** BV_645_/BV_395_ representing the degree of bone mineralization. Control mice: n = 7, Loaded mice: n = 7. Significance criteria: # represents comparisons between Young and Aged mice at each timepoint (# p < 0.05, ## p < 0.01, ### p < 0.001, #### p < 0.0001), * represents comparisons within groups between timepoints (* p < 0.05, ** p < 0.01, *** p < 0.001, **** p < 0.0001).

In DC regions, BV/TV diverged between Control and Loaded groups from week 4 onwards with significant differences at week 5 (p ≤ 0.01), week 6 (p ≤ 0.001) and week 7 (p ≤ 0.001) (Fig 6. [A]). At week 7, BV/TV in the DC region was 35.0 ± 6.2% in Control mice versus 66.4 ± 11.2% in Loaded mice (p ≤ 0.001). Dynamic bone morphometric parameters were consistent with a strong anabolic response to mechanical loading in the DC regions of Aged mice. In both Control and Loaded groups, BFR and BRR declined between weeks 4 – 7 (Fig 6. [E]); however, mechanical loading resulted in a net bone formation response with a sustained BFR (significantly higher at week 6 and week 7; p ≤ 0.05) and a diminished BRR (significantly lower at week 7; p ≤ 0.05) (Fig 6. [E]). Progressive (and significant) increases in the degree of mineralization were observed in the DC region of both Control and Loaded groups between weeks 4 – 7 (Fig 6. [I]). At week 7, the fraction of mineralized bone was significantly greater in Loaded mice compared to Control mice (p ≤ 0.05) (Fig 6. [I]).

In contrast to the pronounced response observed in DC regions, peripheral regions (DP, FP) exhibited a limited response to mechanical loading (Fig 6.). Non-significant trends in BV/TV towards gradual increases in Loaded mice and decreases in Control mice were observed between weeks 4 – 7 (Fig 6. [B][D]). Dynamic bone morphometric parameters in peripheral regions suggested a delayed response to mechanical loading. Non-significant but distinct trends in BFR were present between weeks 5 – 7, characterized by gradual increases in Loaded mice and declines in Control mice (Fig 6. [F][H]). In response to mechanical loading, BRR in peripheral regions was significantly suppressed between weeks 4 – 5 (FP: p ≤ 0.05), weeks 5 – 6 (DP: p ≤ 0.05, FP: p ≤ 0.01) and weeks 6 – 7 (DP: p ≤ 0.05, FP: p ≤ 0.01) (Fig 6. [F][H]).

Collectively, these findings demonstrate that mechanosensitivity of the regenerative response is retained in Aged PolgA mice; however, in contrast to Young mice, the anabolic response is localized to the defect center and limited in peripheral regions.

## Discussion

Although the mechanical environment is of critical importance to the regenerative response in bone, there is no consensus on the effects of aging on the mechanosensitivity of bone regeneration. Using an established femur-defect model in PolgA mice, we establish the PolgA mouse strain as a relevant preclinical model in investigations of age-dependent mechanobiological responses in bone regeneration. Aged PolgA mice were found to recapitulate key characteristics of age-related impairments in bone regeneration, including delayed fracture healing, impaired bone formation and resorption dynamics, attenuated osteogenesis, and delayed mineralization of newly formed bone. Notably, the regenerative response in Aged mice was found to differ both spatially and temporally from Young mice, with delayed bone formation in the defect center and reduced osteogenic activity in peripheral regions. We demonstrated that cyclic loading significantly enhanced regeneration in Young PolgA mice, and this mechano-responsiveness was remarkably retained in Aged PolgA mice. Furthermore, age-related shifts in baseline regeneration likely contributed to the spatial localization and temporal delay in the anabolic response to cyclic mechanical loading in Aged bone. These findings provide direct evidence that aging alters the temporal and spatial dynamics of bone regeneration without abolishing the capacity of bone to respond to mechanical stimuli. Our findings thus suggest that mechanical stimulation therapies have the potential to be promising interventions in the treatment of challenging fractures in elderly patients. Ongoing analyses apply our recently established “spatial mechanomics” approach (Fig 7.) to further explore age-dependent mechanobiological mechanisms in bone regeneration and to identify novel mechanoresponsive targets and strategies to enhance repair in compromised healing environments.

**Fig. 7.**
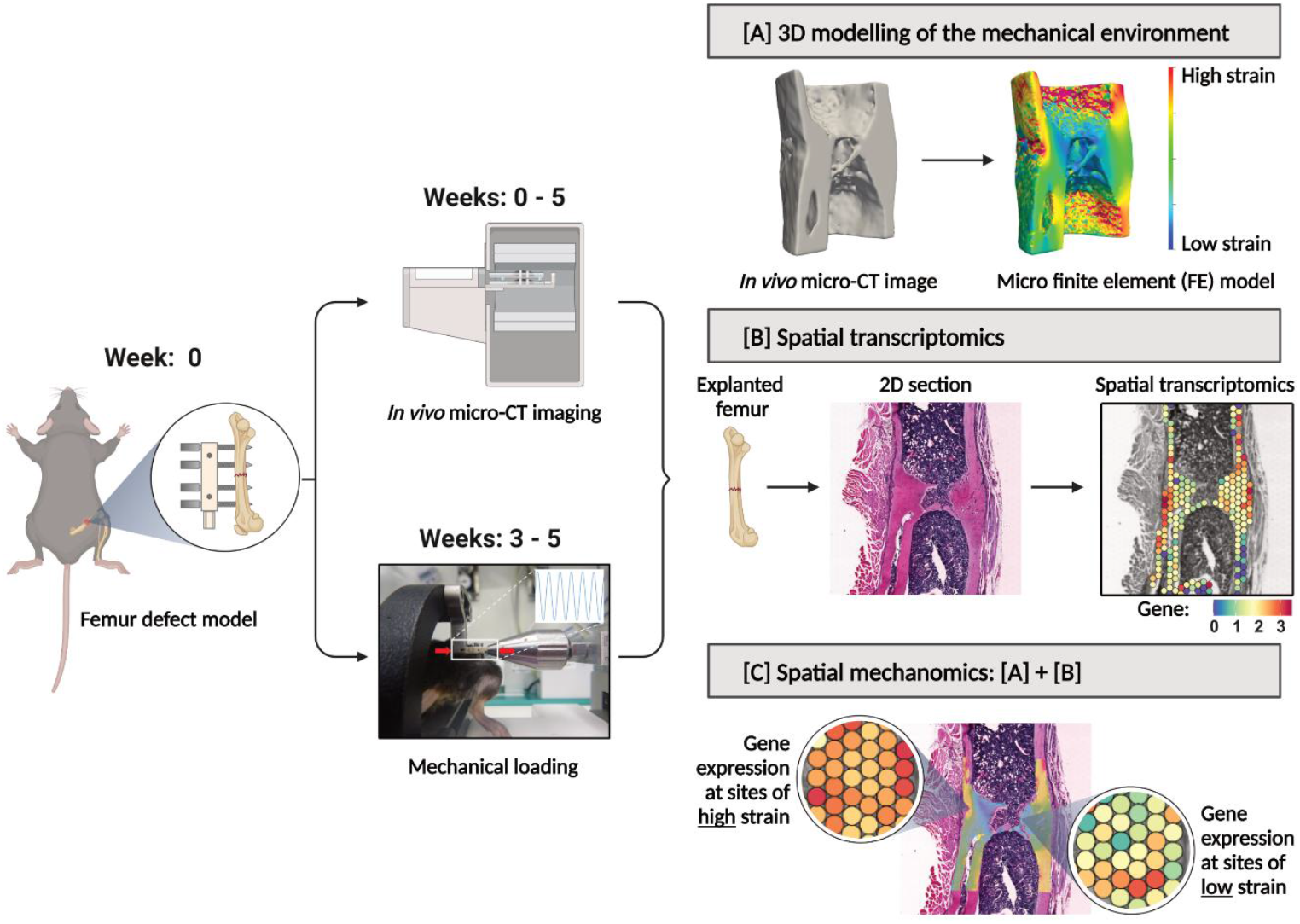
Overview of our “spatial mechanomics” approach. Our approach permits the generation and spatial integration of multimodal datasets (CT bone morphology data, 3D mechanical environments, and spatially resolved gene expression data) from a single fracture site. Illustration reproduced from Mathavan et al.^11^.

## Materials and Methods

### Experimental Design

Our objectives were to investigate the influence of aging on the mechano-responsiveness of the regenerative response in bone and to investigate the efficacy of cyclic mechanical loading in harnessing the mechanosensitivity of aged bone. As illustrated in Fig. 1, our experimental design consists of experiments performed using homozygous PolgA^D257A/D257A^mice – a mouse model of premature aging^15,21^. Using an established mouse femur defect model^12^, all mice received femoral osteotomies in their right femur. *In vivo* micro-CT imaging of the fracture site was subsequently performed each week in all mice. Mice which exhibited bridging of the fracture site at 3 weeks after surgery were subdivided into Control or Loaded groups and received either sham-loading or cyclic mechanical loading respectively.

### Ethics statement

All mouse experiments were performed in accordance with relevant national regulations (Swiss Animal Welfare Act, TSchG, and Swiss Animal Welfare Ordinance, TSchV) and authorized by the Zürich Cantonal Veterinary Office (approved licenses: ZH229/2019 and ZH033/2022; Kantonales Veterinäramt Zürich, Zurich, Switzerland).

### Mouse line

All PolgA^D257A/D257A^mice were bred, monitored, and maintained under specific pathogen–free conditions at the ETH Phenomics Centre, ETH Zürich (12-hour light/12-hour dark cycle, ad libitum access to maintenance feed and water).

### Mouse femur defect model

12-week-old or “Young” female PolgA mice (n = 15, age = 12.6 ± 0.6 weeks) and 35-week-old or “Aged” female PolgA mice (n = 14, age = 35.3 ± 0.2 weeks) received a mid-diaphyseal femoral fracture (defect sizes in young mice: 0.57 ± 0.1 mm, defect sizes in aged mice: 0.42 ± 0.1 mm) stabilized with an external fixator (license: ZH229/19, ZH033/2022 Kantonales Veterinäramt Zürich, Switzerland). Femoral fractures were introduced using an established osteotomy surgery protocol^12^. In all mice, an external fixator (Mouse ExFix, RISystem, Davos, Switzerland) was first positioned at the craniolateral aspect of the right femur using four mounting pins. Osteotomies were subsequently performed using a 0.66 mm or 0.44 mm Gigli wire saw. Peri-operative analgesia (25 mg/liter, Tramal, Gruenenthal GmbH, Aachen, Germany) was provided via the drinking water 2 days before surgery until the third post-operative day. Anesthesia for all animal procedures (surgery, *in vivo* imaging, and mechanical loading) was administered using isoflurane (induction/maintenance: 3%, 1.5% to 2.5% isoflurane/oxygen).

### In vivo micro-CT imaging

*In vivo* micro-CT imaging of the fracture site was performed weekly in all mice (weeks 0 to 7; vivaCT 80, Scanco Medical AG, Brüttisellen, Switzerland) (10.5 μm nominal resolution, 55 kVp, 145 μA, 350 ms integration time, 500 projections per 180°, 21 mm field of view, scan duration ca. 15 min). To avoid motion artifacts during scanning, a custom-designed holder was used to secure the external fixator and the volume between the two inner screws of the external fixator encompassing the fracture site and adjacent cortices was scanned. Registration of time-lapsed *in vivo* images permits visualization of sites of bone formation, quiescence, and resorption^22^. Reconstructed images from all timepoints were sequentially registered for each mouse, as described previously^23^. Images were Gaussian filtered (sigma 1.2, support 1) and bone volumes (BVs) were computed (threshold: 395 mg HA/cm^3^) in four non-overlapping volumes of interest (VOIs): the defect center (DC), the defect periphery (DP), the existing fracture cortices together with the medullary cavity (FC), and the cortex periphery (FP) (Fig. 3)^23^. Bone morphometric indices (bone volume fraction: bone volume BV/ total volume TV, bone formation rate: BFR, bone resorption rate: BRR) were evaluated within each VOI. To quantify mineralization progression, voxels exceeding a second threshold of 645 mg HA/cm^3^were classified as highly mineralized, and the ratio of high to low mineralized tissue (BV_645_/BV_395_) was calculated. Calculated parameters per VOI were normalized with respect to the central VOIs (DC and FC) which represent the total volume (TV) of intact bone^23^: thus, DC/DC, DP/DC, FC/FC, and FP/FC. Defect sizes (h) were calculated using the following formula: h =2 DC/(CSA_P + CSA_D), where DC is the defect volume at week 0 and CSA_P and CSA_D represent the proximal and distal cross-sectional areas, respectively, which are situated directly adjacent to the fracture site.

### In vivo cyclic mechanical loading

In 20-week-old C57BL/6J mice, we have demonstrated that our femur defect model results in bridging of the fracture site at three weeks post-surgery in all mice^12^. In Young and Aged PolgA mice, the femoral defect model resulted in variable healing outcomes indicative of the impairment in the regenerative response with aging. As a measure of healing progress, fractures sites were classified as (i) unions – if sites were bridged by week 03 post-surgery, (ii) delayed unions – if sites were bridged between weeks 04 – 07 post-surgery, and (iii) non-unions – if no bridging present up to 7 weeks post-surgery. At week 03 post-surgery, fracture sites exhibiting bridging were subdivided into Control and Loaded groups. Control mice (Young: n = 8, Aged: n = 7) received sham loading (0 N, 3x/week). Loaded mice (Young: n = 7, Aged: n = 7) received individualized cyclic mechanical loading (2-16 N, 10Hz, 3000cycles; 3x/week) via the external fixator. Detailed descriptions of the protocols can be found in the literature^12,13^.

The magnitude of the mechanical loading applied to the fracture site at each timepoint was determined by computing the strain distribution within the fracture site of each mouse and then scaling of the strain distribution to achieve a predefined median target strain. The procedure was implemented each week in “real time” within a single anaesthetic session. In short, the mouse was anaesthetized and *in vivo* micro-CT imaging of the fracture site performed. Within the same anaesthetic session, the CT data was reconstructed, processed and a high-resolution *in silico* micro-finite element (micro-FE) model of the fracture site generated. The micro-FE model simulates axial compression by applying a compressive displacement of 1% to determine the mechanical environment at the fracture site. Micro-FE model simulations were computed using ParOSol^24^, a linear micro-FE solver, on an internal computer cluster [48-core Intel(R) Xeon(R) Platinum 8168 CPU @ 2.70 GHz]. Subsequently, the magnitude of the applied load is determined by scaling the strain distribution at the fracture site to achieve a pre-defined median target strain. With this adaptive loading approach, loading is thus individualized to each mouse such that the median induced strain across all loaded mice is of comparable magnitude^13^. Furthermore, to assess whether the applied loading poses a structural failure risk at the fracture site, the simulation identifies sites of high strain (defined by voxels that exceed more than 10,000 με)^13^. If more than 100 voxels exceed this high strain threshold, the loading value is downscaled by 1 N. The structural failure risk analysis is then repeated iteratively until the number of voxels exceeding 10,000 με falls below 100.

### Statistical analysis

Statistical analyses of repeated measurements were performed using two-way ANOVA with Geisser-Greenhouse correction and Bonferroni post hoc testing. Statistical analyses of data derived from single timepoints were assessed for normality (Shapiro–Wilk-Test) and analysed using either an unpaired two-tailed Student’s t-test or the Mann–Whitney U test, as appropriate. All analyses were conducted using GraphPad Prism 8. The level of statistical significance was set at p < 0.05.

## Acknowledgments

*In vivo* experiments were performed at the ETH Phenomics Center (ETH) of ETH Zürich. Spatial transcriptomics was performed at the Functional Genomics Center Zürich (FGCZ) of the University of Zürich and ETH Zürich. Tissue processing and paraffin embedding were performed at ScopeM at ETH Zürich. Sectioning, staining, and imaging were performed at ScopeM at ETH Zürich, at the Center for Microscopy and Image Analysis (ZMB) at the University of Zürich and at the Institute of Pathology and Molecular Pathology at the Universitäts Spital Zürich (USZ). The spatial transcriptomics workflow was also supported by the Genetic Diversity Centre (GDC) at ETH Zürich.

## Funding

This work was supported by:

European Research Council - Horizon 2020 grant ERC-2016-ADG-741883

European Union Marie Skłodowska-Curie Actions - Horizon 2020 grant 101029062

MechanoHealing-MSCA-IF-2020

Swiss National Science Foundation (IZCOZ0_198152/1, COST Action GEMSTONE)

## Author contributions

Conceptualization: NM, EW, RM

Methodology: NM, GP, EW, RM

Investigation: NM, FM, GP, SL, SW, DY, GK EW

Analysis: NM

Supervision: NM, EW, RM

Visualization: NM

Writing—original draft: NM

Writing—review & editing: All.

## Data and materials availability

All data needed to evaluate the conclusions in the paper are present in the paper and/or the Supplementary Materials.

